# Lunar Reproductive Timing Acts as a Magic Trait and May Recruit Additional Isolating Barriers in Sympatric Marine Midge Populations

**DOI:** 10.64898/2026.09.11.750828

**Authors:** Alexander G.G. Jacobsen, Runa K. Ekrem, Tobias S. Kaiser

## Abstract

Population divergence with gene flow has proven more common than expected, but the underlying mechanisms are not fully understood. Magic traits (i.e. characters involved in both ecological adaptation and assortative mating) offer a tantalizing mechanism to explain the phenomenon, yet empirical demonstrations are rare. Moreover, speciation is assumed to involve genomic processes that couple multiple reproductive barriers, rather than being driven by a single barrier, and it remains unclear how magic traits interact with these processes. Here, we investigate the marine midge *Clunio marinus*, whose reproduction is timed to the extreme spring tide low tides during full moon or new moon. In Roscoff (Brittany, France) there are two sympatric chronotypes, i.e. subpopulations which reproduce only during full or new moons. Lunar reproductive timing is hypothesized to act as a magic trait, as it is both ecologically relevant and isolates reproduction in time. Based on wild-caught mating pairs and laboratory crosses, we show that lunar timing indeed is the major reproductive barrier (RI > 0.87). We also identify additional barriers which further increase reproductive isolation (total RI > 0.94). To explore the genomic basis of these barriers, we perform association testing on several high-F_ST_ loci. Two loci were associated with multiple barriers, and patterns of linkage disequilibrium suggest that these effects may arise from a combination of genetic linkage and pleiotropy. Together, our results indicate that lunar timing played a pivotal role in chronotype divergence, not only as a magic trait but also by promoting the recruitment of additional isolating barriers. We hypothesize that this recruitment is due to the polygenic and complex genetic architecture of lunar timing, highlighting the joint role of magic traits and genomic architecture in divergence with gene flow.

## 1 Introduction

Reproductive isolation (RI) is a defining feature of the biological species concept, making the traits that generate it central to understanding how new species arise. However, the way in which these reproductive barriers form and precipitate genome divergence at the onset of speciation, especially in the presence of gene flow, is still an open question. Theory suggests that speciation requires three elements: a source of divergent selection, a mechanism of RI, and a way to transmit the force of selection to the isolating mechanism (Kirkpatrick & Ravigne, 2002; Nosil, 2012). While the first two ingredients have been widely documented, the third—how selection becomes coupled to isolation—remains the major hurdle for demonstrating sympatric speciation. This difficulty arises because, in the face of gene flow, recombination tends to break apart associations between ecological and mating traits, preventing divergent selection from translating into RI and hindering speciation (Smadja & Butlin, 2011). Magic traits, also known as multiple-effect traits (Smadja & Butlin, 2011), are one solution to this problem. These traits are involved in both the ecology and mating of an organism, whereby divergent ecological selection results in the concomitant divergence of a mating character via pleiotropy (Servedio et al., 2011). Association of the two traits therefore cannot be broken by the shuffling of co-adapted alleles due to recombination, as ecology and mating traits stem from the same locus.

While initially proposed as a mathematical convenience in speciation models, magic traits are hypothesized to be common in nature and potentially quite important for speciation (Servedio et al., 2011). Empirical examples include wing patterns in Heliconius butterflies (Kronforst et al., 2006), aposematic coloration in poison dart frogs (Noonan & Comeault, 2009; Reynolds & Fitzpatrick, 2007), electrical pulses in African weakly electric fish (Feulner et al., 2009), and color mimicry in coral reef fish (Puebla et al., 2007). Although magic traits are often exemplified by mate-signaling/preference traits, any trait that causes individuals to mate in different times or places can play the same role. Traits such as habitat preference or reproductive timing are considered “automatic magic traits” because divergence in the trait automatically creates assortative mating, making them especially potent for facilitating speciation (Johnson et al., 1996; Servedio et al., 2011). Timed reproduction, in particular, is remarkably widespread across the tree of life: plants often flower in distinct seasons (Samach & Coupland, 2000), and many marine organisms restrict spawning to specific days of the lunar cycle (Häfker & Tessmar-Raible, 2020; Kaiser & Neumann, 2021; Raible et al., 2017; Tessmar-Raible et al., 2011). If reproductive timing differences were to operate as a magic trait, they could facilitate speciation in an exceptionally broad array of taxa. Yet despite this potential, there remains limited empirical information for how timed reproduction might induce divergence in natural populations.

The marine midge *Clunio marinus* offers an excellent system to study the role of magic traits at the inception of speciation with gene flow. Occupying rocky intertidal zones along the European Atlantic coast, C. marinus is most notable for timing its development and reproduction to the lunar cycle; more specifically, to the spring tides, i.e. the high amplitude tides that occur around full or new moons (Kaiser, 2014). During spring tide low tides, the short-lived adults (approx. 2 hours) copulate and oviposit on the exposed macroalgae near the water’s surface before perishing as the tide returns. Since the height of low tide changes throughout the month and mating for C. marinus is restricted to the water line during low tide, the depth at which oviposition occurs (and where larvae subsequently live) is determined by the day in which the parents emerge, i.e. their lunar timing (Ekrem et al., 2025; Jacobsen et al., 2026). Competition for space among larvae is hypothesized to drive divergence in lunar timing, as parents emerging away from the spring tide place their offspring at habitat depths with lower larval density (Ekrem et al., 2025; Jacobsen et al., 2026). The resulting difference in reproductive timing would then restrict gene flow (Jacobsen et al., 2026), rendering C. marinus’ lunar emergence a magic trait.

Indeed, in the intertidal zone along the Brittany coast of France, multiple sympatric chronotypes have been found to co-exist (Kaiser et al., 2021). A previous study on the chronotypes in Roscoff tracked the emergence of adults over the lunar month and the distribution of chronotypes in the intertidal zone, and found the exact pattern of shifted emergence and larval depth expected by the aforementioned mechanism, strongly suggesting that C. marinus’ circalunar timing acted as a magic trait (Ekrem et al., 2025). However, despite the difference in lunar timing, these chronotypes show little genome-wide differentiation (Briševac, Peralta, et al., 2023; Kaiser et al., 2021), indicating that divergence is very recent, gene flow is still ongoing, or both. As this previous study only classified chronotypes based on their emergence timing rather than their genotype/ancestry, there may have been a large degree of unobserved “migration in time” that would explain this low genetic divergence. Assessing if lunar timing effectively reduces gene flow, and thus acts as a magic trait, would therefore require quantifying the level of migration between chronotypes.

Moreover, it remains unknown if lunar timing is the only barrier acting to reduce geneflow between chronotypes. Total RI is rarely the product of a single barrier, but rather is produced by the joint action of several barriers (Butlin & Smadja, 2018). Slight differences in diel emergence timing (i.e. the time of day individuals emerge) have been observed for these chronotypes (Kaiser et al. 2021, Ekrem, Jacobsen et al. 2025), but it is unclear to what degree these differences might also contribute to RI. Other barriers operating at other life history stages, such as gametic incompatibility or hybrid depression, may also contribute to RI, but they have yet to be tested.

Finally, if there are other barriers to reproduction, an important question is whether these additional barriers share a common genetic basis with lunar timing, or whether they are independent. Barriers that are coupled, either through physical linkage or pleiotropy, can reinforce one another, producing a more cohesive overall barrier to gene flow than would arise if the same barriers were distributed independently across the genome and thus more readily uncoupled by recombination (Butlin & Smadja, 2018; Smadja & Butlin, 2011). If lunar timing shares a genetic basis with additional barriers, this may have interesting evolutionary implications for its role as a magic trait, suggesting that lunar timing was not only important for reducing gene flow, but also for recruiting additional “non-magic” barriers via a shared or linked genetic basis.

Our study has three aims. First, we asses if lunar timing acts truly as a magic trait by quantifying its contribution towards reproductive isolation for *C. marinus* chronotypes. Second, we test for other barriers to reproduction to determine if lunar timing acts in isolation. Third, if there are other barriers acting, we then investigate their genetic basis to understand how much is shared with lunar timing.

We focus on the two sympatric chronotypes found in Roscoff: a full-moon chronotype (Ros-2FM, hereafter FM) and a new-moon chronotype (Ros-2NM, hereafter NM). Using a dataset of mating pairs collected in the field and subsequently genotyped at high-F_ST_ loci, together with a series of controlled laboratory crosses and emergence data under common-garden laboratory conditions, we identify reproductive barriers acting between chronotypes across their life cycle. We then use these complementary datasets to quantify the absolute strength of each barrier and estimate the total RI between the chronotypes. Finally, by testing for associations between barrier traits and genotypes at the high-F_ST_ loci we evaluate which genomic regions contribute to each barrier, then by calculating linkage disequilibrium around each associated locus we assess the potential role physical linkage and pleiotropy in facilitating divergence. Together, these results clarify the central role of reproductive timing—and magic traits more broadly—in the buildup of RI and illustrate how genomic architecture can shape the trajectory of speciation.

## 2 Methods

### 2.1 Study system

C. marinus occupies rocky stretches of the European Atlantic coast, where individuals spend most of their lives as larvae, feeding in the macroalgae turfs that grow in the bottom of the intertidal zone. In the days surrounding the spring tides, adults emerge when the receding tide exposes the larval substrates. The winged and motile males scan the surface of the water for the wingless and immotile females, and once finding the female, he will copulate with her and then take her to a near-by patch of exposed substrate so that she can oviposit. Adults typically live no longer than 2 hours, so copulation and oviposition occur within a single low tide.

### 2.2 Field Sampling of copulae and genotyping

Sampling: To assess barriers to reproduction for the chronotypes in the wild, we sampled copulating individuals (i.e. female and male pairs actively reproducing) in the field in Roscoff (48◦43’ 39.95” N, 3◦59’ 4.08” E) from March-May of 2019, encompassing two full moon and two new moon emergence peaks. We collected copulae in late afternoon and evenings and recorded the time of capture in 30-minute intervals. Intervals were also recorded when no emerging adults were found. Sampling was accomplished by either visually scanning the water’s surface for adults and then catching them in a sieve, or during periods of high emergence, sweeping the water’s surface with the sieve near patches of algae. We placed captured copulae in petri dishes and allowed them to finish mating and ovipositing. The adults were preserved in ethanol and stored at -20◦C. Eggs were placed in a fridge at 4◦C for 24 hours to delay development before being shipped to the laboratory. These eggs were hatched, and offspring were reared and phenotyped under common garden conditions (Supplemental methods) for use in further analyses described below. In total, we collected 420 couples.

The lunar day of each individual was assigned based on the lunar day of the low tide in which it emerged. The date of the first low tide during sampling was converted to lunar day using the pyephem library (Rhodes, 2020) in python 3.9.13 (Van Rossum & Drake, 1995), and every other low tide was then considered a subsequent lunar day. If sampling occurred across midnight, the time of emergence was converted to the number of hours after midnight on the previous lunar day to avoid discontinuity (when the time changes from 24:00 to 00:00).

Genotyping: Though chronotypes are generally defined by their emergence timing, individuals can sometimes emerge at the “wrong time” (Briševac, Peralta, et al., 2023), i.e. on a day that does not reflect their ancestry. Therefore, to infer the ancestry of individuals, we employed a genotyping-by-sequencing approach, targeting 11 of the most genetically diverged loci between FM and NM, as identified by an F_ST_ scan of 24 full genomes from either chronotype (Briševac, Peralta, et al., 2023). A version of this genotyping approach is also described in (Ekrem et al., 2025).

In brief, we designed primers (Supplemental Table 1) to amplify a small region around the most diverged SNP in each locus (i.e. F_ST_ peak) to use as an ancestrally informative marker. DNA from entire, macerated individuals was extracted following (Reineke et al., 1998) and the selected regions were PCR amplified (for conditions and procedure see Ekrem, Jacobsen et al. 2025). We then sequenced the amplicons, aligned them to the C. marinus reference genome, and called SNPs using GATK (Poplin et al., 2017). Additionally, locus 3 (the *period* locus) contained an insertion-deletion (indel; approx. 600 bp) associated with chronotype, which caused inconsistent sequencing. We therefore designed new primers (Supplemental Table 1) to amplify across this indel in order to score the locus in an agarose gel, using the presence of the indel as the marker for this locus. We applied the maximum likelihood genotyping approach described in (Buerkle, 2005) to assign an ancestry proportion to individuals, using the 24 full genomes for FM and NM to calculate the allele frequencies of the parental populations. We were able to genotype 797 individuals from 387 couples.

### 2.4 Crossing chronotypes in the laboratory

While our field couple data represented the mating patterns of the chronotypes in the wild, it contained comparatively few heterospecific crosses where we would expect to see the greatest effect of barriers to reproduction. Therefore, we complimented our field data with controlled crosses between chronotypes in the laboratory. We raised NM and FM individuals in climate-controlled culture rooms under identical conditions (Supplemental Methods). However, we shifted the moonlight and light-dark cycles for each chronotype to synchronize their emergence to the same days of the month as well as the same hours of the day, thereby largely obliterating the effects of reproductive timing and making other barriers more readily visible. Crossing was done over three separate emergence peaks spanning three months.

During days of emergence, boxes were brought to a common laboratory shortly before emergence was expected to start and any already emerged adults were cleared from the boxes. Newly emerging pupae were captured as soon as they rose to the water surface and were allowed to finish eclosing individually in a small petri dish with some seawater. The chronotype, sex, and time were recorded for each individual. We crossed individuals in small (8x8 cm) sterile acrylic boxes, with a shallow layer of sterile 50:50 diH_2_O:seawater on the bottom. The female was first added into the box, followed by the male.

As soon as the male entered the box a timer was started and the cross was observed. Once a copula was formed, as determined by the individuals being locked together end-to-end by their genitalia and freely moving, the timer was stopped. This time was recorded, as well as the time (hour and minute) that the individuals were added to the crossing box. If the individuals did not form a copula after 10 minutes, observation was stopped and a value of 10:00+ was recorded as the time to copulation. Boxes were placed in a culture room overnight to allow completion of copulation and oviposition. The following day, any eggs found in the crossing boxes were placed in a petri dish with sterile 50:50 diH_2_O:seawater. Eggs were then left to develop at 19°C for 4 days under a light-dark cycle with 16 hours of light and 8 hours of darkness (LD 16:8). On day 5, the number of eggs, as well as how many were fertilized (i.e. had developed into an embryo) were counted. The eggs were then left for an additional 5-10 days under the same conditions to allow the embryos to develop into larvae. We then counted the number of larvae that had successfully hatched from the egg clutch. In total, we performed 240 crosses.

### 3.4 Testing for barriers

The lunar and circadian timing of emergence was previously characterized for FM and NM in laboratory conditions (Kaiser et al., 2021). To assess timing difference in the field, we visualized the distribution of emergence for chronotypes for the field data over the sampling period by plotting the joint distribution of emergence day and emergence timing, coloring points by ancestry proportion (and not by phenotype as in Ekrem, Jacobsen et al. 2025). The emergence distribution over lunar days as well as over hours within the day were plotted in the margins.

Using our complementary field and laboratory data sets we could also test for the presence of additional reproductive barriers at nearly all stages of C. marinus’ life cycle. This includes a barrier outside of timing preventing heterospecific mating (i.e. mate preference), a barrier preventing heterospecific fertilization (i.e. gametic incompatibility), a barrier reducing the hatching rate or survival of hybrid offspring (i.e. hybrid depression), and finally a barrier preventing or reducing the number of offspring from hybrid individuals (i.e. hybrid infertility).

#### 3.4.1 Mate preference

To test for mate preference in our field data, we performed a permutation test using the Spearman correlation coefficient between male and female ancestry proportions over couples as the test statistic. We shuffled males between couples within each time-point (i.e. 30-minute sampling interval per lunar day) 1000 times, calculating the Spearman correlation each time to generate a null distribution for ancestry correlation without mate preference. Significance was checked by comparing the observed correlation coefficient to the null distribution. To avoid issues with sample size, data were subset to include only time-points with a minimum of 4 couples.

We also tested for evidence of mate-preference in the laboratory crosses. If males preferred conspecifics, then they may be reluctant to copulate with a heterospecific female, translating into a longer time for copulation or no copulation at all. To test for this, we used a Cox Proportional Hazards model from the lifelines package (Davidson-Pilon, 2019) in python with the formula crossing time ∼ cross type ∗ female age + number of eggs. Cross type corresponded to crosses within or between chronotypes, female age was the difference between the time that the female emerged from the water and the time the cross was initiated, and the number of eggs was the number of eggs produced by the female. Number of eggs was included in the model as its addition improved the model’s fit, as noted through a lower AIC.

#### 3.4.2 Gametic incompatibility and hybrid infertility

We used the fertilization rate of our field couples to simultaneously test for gametic incompatibility and hybrid infertility. Three things could affect the fertilization rate: gametic incompatibility due to a difference in parent ancestry, the heterozygosity of the mother reducing her fertility (due to hybrid infertility), or heterozygosity in the father reducing his fertility (also due to hybrid infertility). To test for these effects, and given that the number of fertilized eggs was zero inflated, we used a zero-altered negative binomial (ZANB) model, modeling both the probability of any eggs in a clutch being fertilized at all, and then the fertilization rate for clutches that were fertilized. We included the ancestry difference between the parents (|*male ancestry* − *female ancestry*|), the female heterozygosity, and the male heterozygosity as predictors. Heterozygosity was taken as the proportion of heterozygous loci out of all loci that were successfully genotyped. We included the log of the total number of eggs a female produced as an offset. The model was fit using the statsmodels package in python (Seabold & Perktold, 2010).

In the laboratory crosses, as we only had purebred individuals available, we could only test for gametic incompatibility. Using the crossing dataset, significant differences in fertilization rate were checked between crossing directions via a Kruzkal Wallace test using the scypi library (Virtanen et al., 2020). A Dunn test was used posthoc to determine which groups were significantly different.

#### 3.4.3 Hybrid depression

To test for hybrid depression in our field couples, we used the survival rate of offspring raised in the laboratory (i.e. the number of offspring that emerged for each cross). Since it was possible for the offspring to reproduce and oviposit before being collected, we only considered offspring emerging in the first two months to avoid including individuals from a possible second generation. We calculated the expected heterozygosity in the offspring based off the genotypes of the parents. To do this, we assumed that each locus segregates in a Mendelian ratio and calculated the proportion of heterozygous offspring at each locus. Averaging across loci gave the expected heterozygosity for offspring from each couple. We then used ZANB model as before to test for an effect of expected heterozygosity on survival rate, including the log of the number of fertilized eggs for the couple as an offset in the negative binomial component of the model. This model was again fit using the statsmodels package.

For the laboratory crosses, we again used a Kruzkal Wallace test followed by a post-hoc Dunn test to check for significant differences in the hatching rate for different crossing directions.

### 3.5 Quantifying the strength of barriers

To quantify the strength of the barriers to reproduction, as well as quantify the total RI between chronotypes, we used the framework from (Sobel & Chen, 2014).

This framework presents a general formula for RI, which is

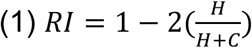

where H is the number of heterospecific matings and C is the number of conspecific matings. Since this framework requires a binary classification to count heterospecific vs conspecific matings, we set a threshold of 0.5 for our ancestry proportions, where individuals above 0.5 were classified as FM and those below 0.5 were classified as NM.

For all measures, confidence intervals were generated by bootstrapping the dataset used for the calculation. In cases where calculations were based on model predictions, the model was refit to the resampled data for each bootstrap.

#### 3.5.1 Lunar and diel timing

To calculate the strength of RI due to lunar or diel timing, we modified equations 4B and 4S1 from (Sobel & Chen, 2014), which are adjusted for spatial/temporal overlap in varying proportions, to calculate *H* and *C* for equation 1. For these calculations, we use both the emergence data for individuals in the field, as well as the common-garden laboratory emergence data from (Kaiser et al., 2021). As the number of individuals collected for FM and NM in the field were not reliable estimates of total abundance, and the total count of individuals per chronotype in the laboratory were not biologically meaningful, we modified these formulas to use proportions of individuals within chronotype rather than counts.

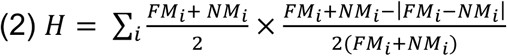

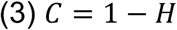

*FM_i_* and *NM_i_* are the proportions of FM and NM individuals present during time-point *i.* The left-hand term in H represents the proportion of all matings that occur during time-point *i* in which heterospecific matings are possible. When chronotypes are present in unequal numbers, heterospecific matings are limited by the less abundant chronotype, such that only the shared fraction of individuals from each chronotype can have the opportunity to mate with the other chronotype. Under random mating, half of these potential pairings are expected to be heterospecific (hence the 2 in the denominator). The left-hand term then scales this expectation by the proportion of matings that occur during time-point *i* to all matings. Because mating frequencies are calculated as proportions of individuals per time-point within each chronotype, and there are two chronotypes, the total number of matings across all time-points sums to 2. Calculations were made using both lunar days and diel time (in 30-minute increments) as the time-points.

#### 3.5.2 Mate preference

To calculate the strength of mate preference, we leveraged the Cox Proportional Hazards model that we had previously fit to the laboratory crosses to predict the proportion of successful matings for con- and heterospecific crosses at the 10-minute mark, which was the extent of the measured time to copulation in our data. We modified the general formula for RI to be

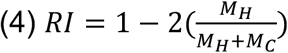

where *M_H_* and *M_C_* were the proportion of hetero- and conspecific matings, respectively. However, in our model there was also a strong effect of the females age on mating probability. Under common-garden conditions, males tend to emerge earlier than females and the NM chronotype tends to emerge earlier in the day than the FM chronotype (Ekrem et al., 2025; Kaiser et al., 2021), meaning that there should be an asymmetry in the age of heterospecific females for crossing directions (FM♂-NM♀ or NM♂-FM♀). We therefore estimated the average expected age of a heterospecific female for each direction by using the distribution of laboratory emergence times for FM and NM males and females. To accomplish this, we generated 1,000,000 random pairs of male and female emergence times and calculated the age of the female at copulation by subtracting the males age from the females, setting negative values (i.e. where the male emerged first) to zero. We additionally censored all pairs with an age difference greater than 1.85 hours (the choice of this threshold is discussed below), then calculated the average age of females for crosses within chronotype and for each crossing direction. We used these ages to calculate model predictions, and subsequently the strength of RI for each crossing direction.

The age threshold of 1.85 hours was chosen for several reasons. In natural settings, adults are thought to survive only during the approximately two-hour window of low tide when the larval habitat is exposed, making larger age differences unlikely. Consistent with this, laboratory observations suggest that adults maintained for longer periods are severely energy-depleted and do not readily mate. 1.85 hours corresponds to the age of the oldest female in the crossing data used to fit the Cox proportional hazards model; including older ages would therefore require extrapolation. However, to better understand the impact that our choice of age threshold had on the calculated values of RI, we performed a sensitivity analysis by calculating the average female age for different crosses, the RI for the mate preference barrier, and the relative RI of the mate preference barrier to the total RI under different age thresholds.

#### 3.5.3 Hybrid infertility

We again modified the general formula for RI to calculate the strength of hybrid fertility to be

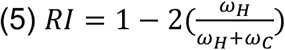

where *ω_H_* was the fitness (i.e. number of fertilized eggs) for hybrid individuals, and *ω_C_* was the fitness of purebred individuals. The number of fertilized eggs for hybrid and purebred individuals was predicted using a ZANB model fit to our data, with only the female’s heterozygosity included as a predictor. Since none of the alleles were fixed within either chronotype, to calculate the expected heterozygosity in a hybrid or purebred individual, we used the following formulas:

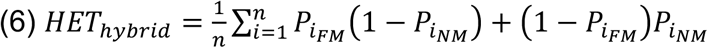

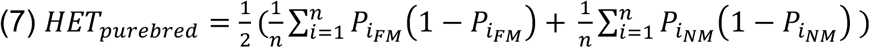

where *P_iFM_* and *P_iNM_* were the minor allele frequencies at the *i*th locus for FM and NM, calculated from the 24 whole genome reference individuals used previously for genotyping. The values for *HE*T_ℎ*ybrid*_ and *HE*T*_purebred_* were then plugged-in to the zero altered negative binomial model to get *ω_H_* and *ω_C_*, respectively.

#### 3.5.4 Total reproductive isolation

To calculate the total reproductive isolation, we modified equation 4A from (Sobel & Chen, 2014):

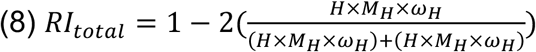

For the calculation of *H* when lunar and diel timing barriers were combined, we used the 30-minute sampling increments within each lunar day as the time-point. Otherwise, all other values were calculated as before for the individual barriers.

For the field data, since we found no evidence of mate preference, we did not include *M_H_* and *M_C_* in the calculation of total RI measured for the field. Likewise, since our data did not allow us to test for hybrid infertility in the laboratory, we did not include *ω_H_* and *ω_C_* in the calculation of total RI in the for the laboratory measurements.

### 3.6 Barrier effects at individual loci

The high-F_ST_ loci that we genotyped in our field couples were not only indicative of ancestry, but also likely represented regions in the genome resistant to gene flow. In other words, these loci potentially contained genes or features involved in the different barriers to reproduction. To get an indication of the genomic architecture of the potential barriers, we therefore inferred the barrier effects of each locus from the genotypes of the field couples.

Through multiple regression, we estimated the effect size of each locus for circalunar emergence timing, circadian emergence timing, and hybrid infertility. Since there was no evidence for mate preference in the field couples, and no evidence at all for gametic incompatibility or hybrid depression, we did not test for these at the locus level. Prior to fitting the regression models, we calculated the variance inflation factors to ensure predictors (i.e. loci genotypes) were not co-linear using the statsmodels package (Seabold & Perktold, 2010). With all tests, we corrected p-values within each testing family using the Benjamini-Hochberg method (Benjamini & Hochberg, 1995).

For circalunar and circadian emergence, we used Cox Proportional Hazards regression after ensuring our data met the assumptions of the model. The formulas used were lunar day ∼ locus1 + locus2. . . locus11 + hour and hour ∼ locus1 + locus2. . . locus11 + lunar day, where lunar day was the day of emergence in the lunar cycle and hour was the time when the individual emerged. Hour and lunar day were included as covariates as these were correlated phenotypes. For hybrid infertility, we used a ZANB model as before, but using the heterozygosity of individual loci for the female in each cross as the predictors. Again, we included the log of the total number of eggs as an offset.

To further investigate the genomic architecture for loci that showed multiple association signals (loci 3 and 8), we visualized the linkage disequilibrium (LD) structure around each locus using 48 whole-genome sequences (24 per chronotype) from (Briševac, Peralta, et al., 2023). LD calculations were done with PLINK (Purcell et al., 2007). To best visualize the physical LD (i.e. LD not from population structure), we calculated LD only within each chronotype. As some of the individuals in (Briševac, Peralta, et al., 2023) might have been hybrids or migrants in time, we used a PCA of unlinked SNPs to exclude individuals that did not cluster neatly with others of their chronotype. This left us with 16 individuals per chronotype for the LD calculations. For each locus, we then selected the 5 closest SNPs to our marker with a minor allele frequency greater than 0.1 to use as proxy SNPs. This was done as our marker SNPs were nearly fixed in one chronotype, making LD calculations unreliable. We calculated the pairwise LD between the proxy SNPs and all other SNPs in a 1MB window. Finally, to plot a smooth curve of LD from each marker combining the information from all proxies, we fit a LOESS to the pairwise LD value for each proxy SNP for each chronotype. This was done separately for values upstream and downstream of the marker SNP, giving us 10 curves per direction. We then plotted the median values of these curves along with the upper and lower quartiles.

## 3 Results

### 3.1 Timing differences in the wild

Under laboratory conditions, the sympatric Roscoff FM and NM chronotypes exhibit marked differences in reproductive timing, both in the day in the month and the hour in the day (Supplemental Figure 1, Kaiser et al., 2021). Field data suggest that these differences are also present in natural conditions (Ekrem et al., 2025), however, the previous study did not assess the genetic identity of emerging individuals. Therefore, a major goal of this study was to determine if genetically defined FM and NM strains align with phenotypically defined FM and NM chronotypes in natural conditions as well. To that end, we caught and genotyped 387 couples actively mating in the wild. Our genotyping revealed that, while many individuals had an ancestry proportion of 0 or 1 (i.e. purebred), a large fraction of individuals (73%) had intermediate ancestry proportions, with 15% of individuals having ancestry between 0.3 and 0.7 (Figure 1C). This indicates that there are two distinct genetic clusters, but there remains a large degree of gene flow or incomplete lineage sorting between them.

**Figure 1.**
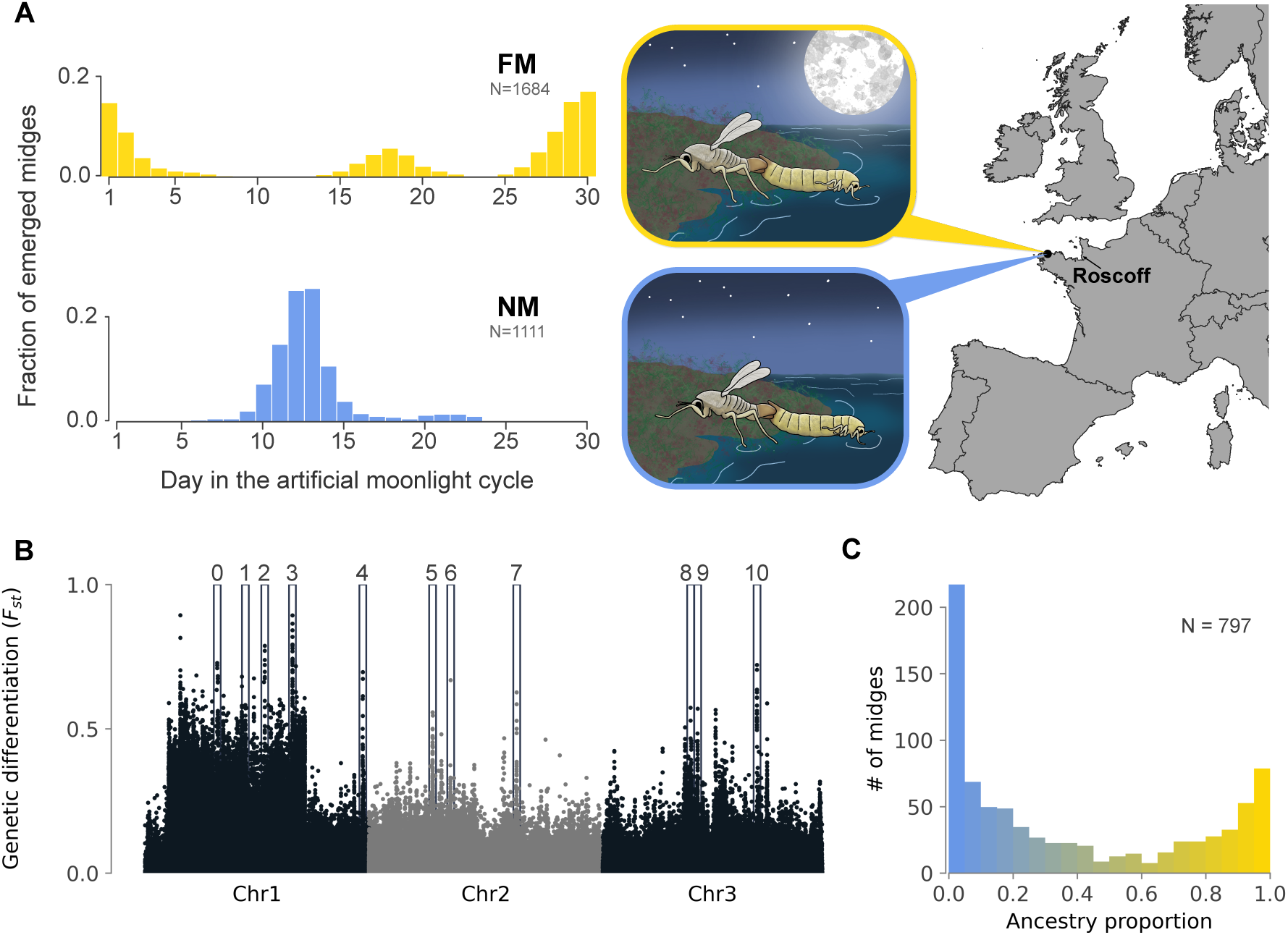
Phenotypes and genotypes of sympatric chronotypes. A) Sympatric chronotypes of C. marinus in Roscoff, France, differ in their lunar emergence timing when kept under common garden conditions in the laboratory, with the FM chronotype (full name: “Ros-2FM”) emerging on the full moon (artificial moonlight day 1) and the NM chronotype (full name: “Ros-2NM”) towards the new moon (artificial moonlight day 15). Small second peaks are laboratory artifacts; data from and more information in Kaiser et al. 2021. B) These chronotypes are genetically diverged (weighted genome-wide F_ST_ = 0.028, Briševac et al., 2023), though differences are primarily restricted to a handful of high-F_ST_ loci across the genome, as well as a large inversion on the first chromosome. Loci used for genotyping individuals in this study are outlined with rectangles and numbered above. C) When genotyping individuals caught in the field, we find many with intermediate ancestry proportions, indicating a large degree of gene flow and/or incomplete lineage sorting.

To test how well phenotypically defined chronotypes and genetically defined strains align, we then plotted the joint lunar/diel distribution of emergence for the field couples (Figure 2A). Emergence of chronotypes in the wild is indeed separated in time, both for the lunar days of emergence as well as the hours of emergence, with lunar emergence peaks approximately 10 days apart and diel emergence peaks roughly 3 hours apart (Figure 2A). These temporal differences correlate largely with genetic ancestry, suggesting phenotypically defined chronotypes and genetically defined strains are the same entities, which for simplicity we only call *chronotypes* from now on. However, the correlation between phenotype and genotype is not perfect and several individuals emerged in the peak contrary to their genotype. This was especially true for genetic FM individuals emerging towards the end of the second NM peak (Figure 2A). Notably, such off-peak emergence is reflected in the FM laboratory strain under certain lighting conditions (Figure 1A, Kaiser et al., 2021). Individuals with hybrid ancestry were generally found between emergence peaks or towards the end of the NM peak.

**Figure 2.**
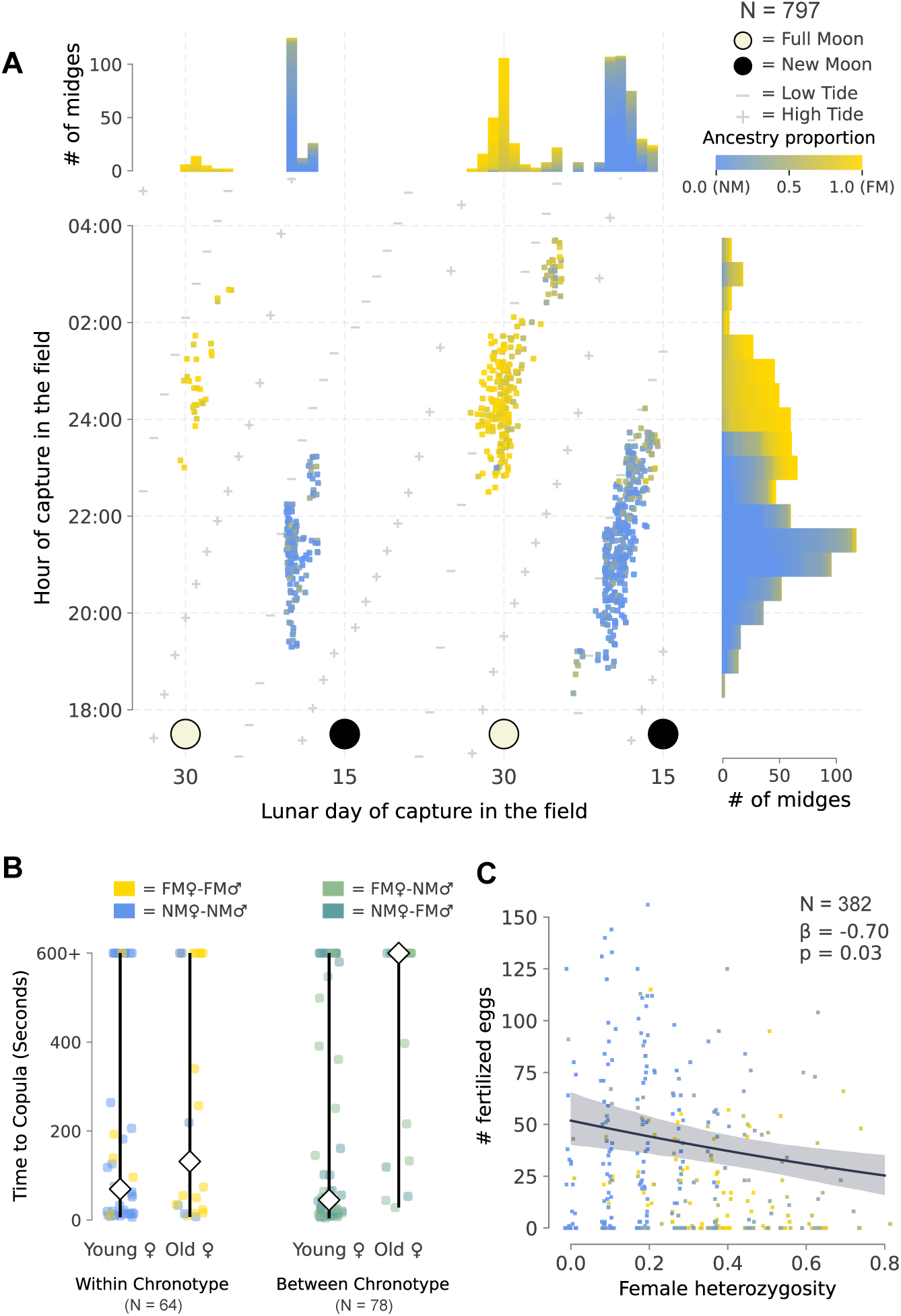
Barriers to reproduction acting between FM and NM chronotypes. A) Individuals (N = 797) were caught while copulating in the field and genotyped to determine their genetic ancestry. Both lunar (top margin) and diel (right margin) emergence timing differences isolate reproduction between chronotypes in the field. Joint lunar and diel emergence timing (middle) isolates reproduction entirely, though occasional “migrants-in-time” can be found, especially towards the ends of the FM or NM peaks. Points are colored according to genetic ancestry. B) When crossed in the laboratory, males will form a copula with a female significantly faster if she is from the same chronotype. This effect is especially apparent when the female is older, i.e. emerged more than 15 minutes prior to crossing. C) For copulae collected in the field, there was a significant effect of female heterozygosity on the fertilization rate of eggs. Heterozygosity was calculated as the proportion of genotyped sites in the female that were heterozygous.

### 4.2 Other barriers to gene flow

The second goal of this study was to determine if timing differences were the only barriers to gene flow, or if there exist other barriers acting after reproductive timing. We tested for 4 additional barriers: mate preference, gametic incompatibility, hybrid depression, and hybrid infertility. All were tested in both laboratory and field crosses except for hybrid infertility, which was only tested in the field data.

#### 4.2.1 Mate preference

By permuting males between couples within a time-point (i.e. 30-minute sampling interval within a lunar day), we found no significant deviation from an assumption of random mating (p = 0.85), indicating no apparent effect of mate preference for our field couples. However, as the separation of mating in time restricts the opportunity for chronotypes to meet, our field data had limited power to detect mate preference: only 8% of time-points contained any couples with an ancestry difference greater that 0.5.

Our controlled crosses in the laboratory afforded much more power to detect mate preference. We timed how long it took males to copulate with females for different crossing types (either within or between chronotype), expecting that if there was mate choice, it would take longer for males to copulate with females of the opposite chronotype. Using Cox’s Proportional Hazards Regression, we found that the crossing type was indeed a significant predictor of the time to copula, in line with our expectation (HR= 1.92, p= 0.02, Figure 2B). Additionally, we found a significant interaction between the crossing type and the age of the female (i.e. the number of minutes since the females emerged before she was placed in the mating arena). The difference in the median time to copula for young and old females was significantly greater for the heterospecific crosses than the conspecific crosses (HR= 0.93, p<0.005, Figure 2B). Notably, many of the heterospecific crosses involving an older female failed to copulate at all (65% for heterospecific crosses compared to 32% for conspecific crosses).

#### 4.2.2 Gametic incompatibility and hybrid infertility

We assessed gametic incompatibility and hybrid infertility in the field couples simultaneously by testing for a significant effect of ancestry difference between parents (gametic incompatibility) or heterozygosity in either parent (hybrid infertility) on the fertilization rate of eggs. We used a ZANB model and found that maternal heterozygosity had a significant effect in the negative binomial component (β=−0.70, p=0.030), i.e. on the number of fertilized eggs. It also had a marginally significant effect in the binomial component of the model (β=−1.20, p = 0.057), i.e. whether any eggs were fertilized vs. not fertilized at all. Neither paternal heterozygosity nor ancestry difference were significant for either component of the hurdle model.

To test if there was any gametic incompatibility between chronotypes for the laboratory crosses, we counted the number of eggs that had been fertilized per cross to calculate the fertilization rate for the different crossing types. Our expectation was that if there was some incompatibility, then the fertilization rate would be lower for the between chronotype crosses. However, when comparing the distribution for fertilization rates using a Kruskal-Wallis test we found no significant difference (H = 6.24, p = 0.10, Supplemental Figure 2A). Taken together with the field data, we can conclude that there is no evidence of gametic incompatibility between chronotypes. However, based on our field data, there appears to be some effect of hybrid infertility, i.e. reduced fertility in females with high heterozygosity.

#### 4.2.3 Hybrid depression

To test for hybrid depression in our field crosses, we again used a ZANB model to test for a significant effect of the expected heterozygosity of offspring on the survival rate of larvae (i.e. the number of adults that emerged from each couple’s eggs). We found no significant effect of expected heterozygosity in either component of the model (Binomial: β = 0.90, p= 0.7; Negative binomial: β = 0.56, p= 0.21).

We tested for hybrid depression in the laboratory crosses by comparing the hatching rate of larvae between crossing directions. We expected that if there was some level of hybrid depression preventing the development of larvae, then we would see a reduction in the hatching rate. The hatching rate was calculated as the proportion of fertilized eggs that had hatched from each cross. We tested for a difference between groups using a Kruskal-Wallis test and found that the hatching rate was significantly different (H = 13.44, p = 0.003, Supplemental Figure 2B). A Dunn test confirmed that the NM female to FM male cross in fact had a significantly higher hatching rate than the FM♀ x FM♂ (q = 0.003) or NM♀ x NM♂ (q = 0.016) crosses. Thus, both field couples and laboratory crosses indicate that there is no hybrid depression preventing the development of larvae. Rather, our results suggest that there was some hybrid vigor granted by crossing chronotypes in the laboratory, potentially due to an amelioration of inbreeding depression in the laboratory cultures.

### 4.3 Strength of barriers

We applied the framework of Sobel & Chen (2014) to our data to quantify the strength of RI from each of the barriers we had found acting between FM and NM chronotypes (lunar timing, diel timing, mate preference, hybrid infertility), as well as the total RI between chronotypes. Calculations were performed separately for laboratory and field data, giving us estimates under both natural and controlled conditions. Additionally, we calculated RI for different crossing directions when data allowed, i.e. FM♀ x NM♂ and NM♀ x FM♂. The values for RI, as well as each barrier’s relative contribution to the total RI between chronotypes are listed in Table 1 and summarized in Figure 3. The total RI between chronotypes based off the field data was measured to be 0.94 (0.91-0.95) in the FM♀ x NM♂ direction, and 0.95 (0.94-0.97) in the NM♀ x FM♂ direction. For the laboratory data, the estimated total RI was 0.93 (0.92-0.95) for FM♀ x NM♂ and 0.93 (0.91-0.96) for NM♀ x FM♂. Note, however, that the estimate of total RI in the laboratory is missing the effect of hybrid infertility, which our experimental design did not allow us to measure.

**Figure 3.**
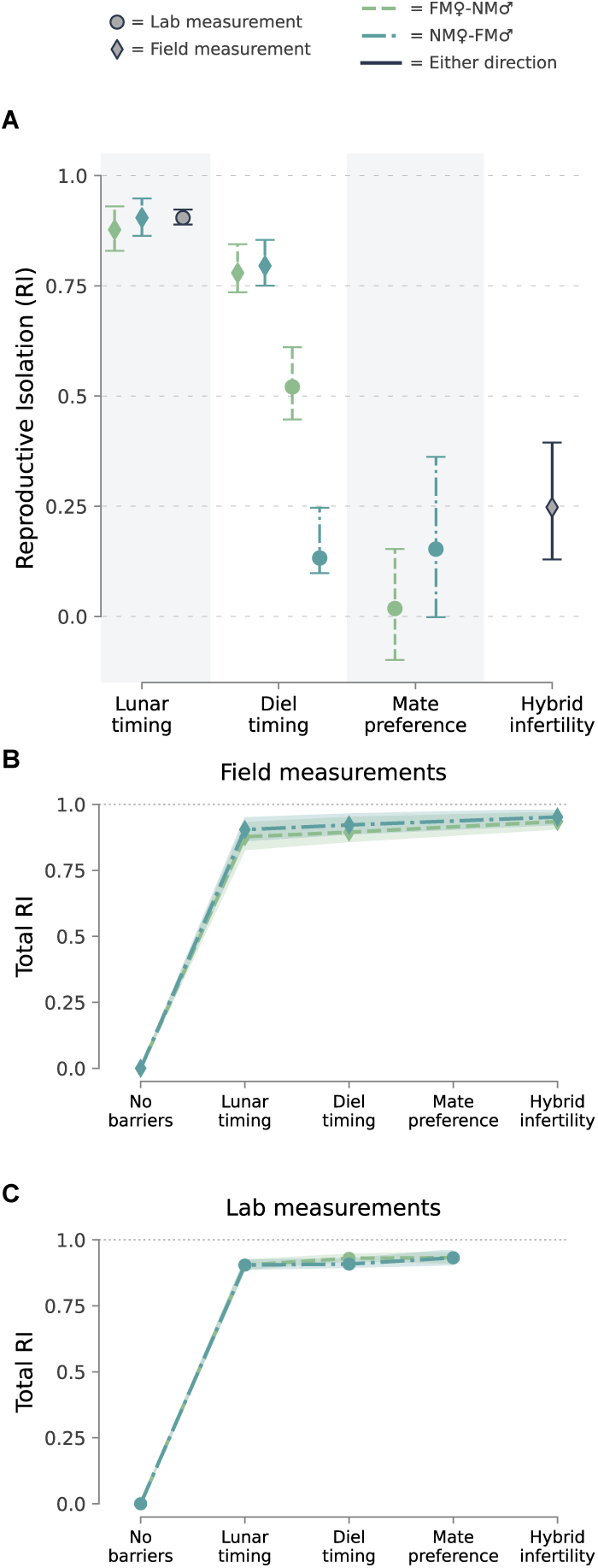
Estimates for reproductive isolation (RI). A) The strengths of barriers to reproduction and (B-C) total reproductive isolation (RI) between FM and NM chronotypes. These values were calculated from measurements made on couples collected in the field (diamonds in panel A and panel B) as well as measurements taken from a set of controlled crosses in the laboratory (circles in panel A and panel C). Values were also calculated separately for each crossing direction (FM♀-NM♂, and NM♀-FM♂). The exact values are listed in Supplemental Tables 1 and 2, and the relative RI values for these barriers are given in table 1.

**Table 1.** The relative strengths of each barrier, calculated for each crossing direction and for measurements taken from the field or laboratory crosses.

| Barrier | Measurement | Direction | Relative RI | 95% CI |
| --- | --- | --- | --- | --- |
| Lunar timing | Field | FM♀-NM♂ | 0.88 | 0.83-0.93 |
|  |  | NM♀-FM♂ | 0.90 | 0.86-0.95 |
|  | Laboratory | FM♀-NM♂ | 0.90 | 0.89-0.92 |
|  |  | NM♀-FM♂ | 0.90 | 0.89-0.92 |
| Diel timing | Field | FM♀-NM♂ | 0.02 | -0.02-0.07 |
|  |  | NM♀-FM♂ | 0.02 | -0.02-0.06 |
|  | Laboratory | FM♀-NM♂ | 0.03 | 0.02-0.04 |
|  |  | NM♀-FM♂ | 0.00 | -0.01-0.02 |
| Mate preference | Laboratory | FM♀-NM♂ | 0.00 | -0.02-0.02 |
|  |  | NM♀-FM♂ | 0.03 | 0.00-0.05 |
| Hybrid infertility | Field | FM♀-NM♂ | 0.04 | 0.02-0.07 |
|  |  | NM♀-FM♂ | 0.03 | 0.01-0.05 |

We found that lunar timing was the barrier with the highest RI, with estimates consistent between the field and laboratory data as well as crossing directions (Figure 3A, Supplemental Table 2). Moreover, lunar timing was by far the barrier with the largest contribution towards the total RI (up to 90%; Table 1, Figure 3 B,C). This indicates not only that lunar timing indeed acted as a magic trait, but was also the most significant barrier to gene flow in the formation of these chronotypes.

Diel timing was the second strongest barrier, with estimates from the field much higher than estimates from the laboratory. The laboratory data also show a large difference in RI between crossing directions, with diel timing posing a stronger barrier in the FM♀ x NM♂ direction than for NM♀ x FM♂. This can be explained in the light of the known sexual dimorphism in adult emergence times: within each strain males emerge on average 1 hour earlier than females (Kaiser et al., 2021). Combined with the diel timing differences between strains, this increases the temporal overlap between the sexes in the NM♀ x FM♂ cross but decreases the overlap in the FM♀ x

NM♂ cross. In either case, in terms of the relative contribution of diel timing to the total RI, diel timing only constitutes a small component.

While our laboratory crosses showed a significant effect of mate preference, the interaction between female age (i.e. minutes after emergence) and the likelihood of copulation within the observation period meant that the value of RI for mate preference was dependent on the female’s age (Supplemental Figure 4A). We therefore estimated the average age of a female for each crossing type, and based on these estimates found the average age of the female in a cross to be 14.9 minutes for conspecific crosses, 10.8 minutes for a FM♀ x NM♂ cross, and 16.6 minutes for a NM♀ x FM♂ cross. Based on these ages, we found that the strength of RI was essentially 0 for the FM♀ x NM♂ direction and small for the NM♀ x FM♂ direction. However, a sensitivity analysis showed that values of RI for mate preference were dependent on our choice of adult lifespan. While RI for the FM♀ x NM♂ direction remained around zero regardless of the age threshold, RI for NM♀ x FM♂ increased with higher age thresholds (Supplemental Figure 4). This, combined with an inability to detect mate preference for the field couples, makes interpretation of the strength of RI for mate preference challenging. Nonetheless, due to its position towards the end in the sequence of reproductive barriers, its contribution to the total RI is small at any age threshold and it likely does not play a major role in reducing gene flow between chronotypes.

Finally, we also found that the value of RI for hybrid infertility was relatively small, though still larger than the estimated RI from mate preference or diel timing in the NM♀ x FM♂ direction (Figure 3A, Supplemental Table 3). Due to its position as the last barrier in the sequence, it only contributes a minor amount to the total RI (Table 1).

### 4.4 The genetic basis of barriers, pleiotropy, and linkage

As the high*-*F_ST_ loci genotyped in our assay may represent barrier loci, we tested the association of barriers with the genotypes at these loci in our field couples, providing insight into the genomic architecture underlying different reproductive barrier traits. To minimize false positives due to population structure, all loci were included simultaneously in each regression model, such that effects were estimated conditional on the genotypes of all other loci. Variance inflation factors were below 3 for all loci, indicating that multicollinearity among predictors was low and that conditional effect estimates were identifiable.

We found that lunar timing was associated with six loci, diel timing with four loci, and hybrid infertility with a single locus (Figure 4A, Table 2). While the loci associated with lunar timing were distributed across the genome, three of the four loci associated with diel timing were located within a previously identified large inversion system on chromosome 1 (Briševac, Peralta, et al., 2023).

**Figure 4.**
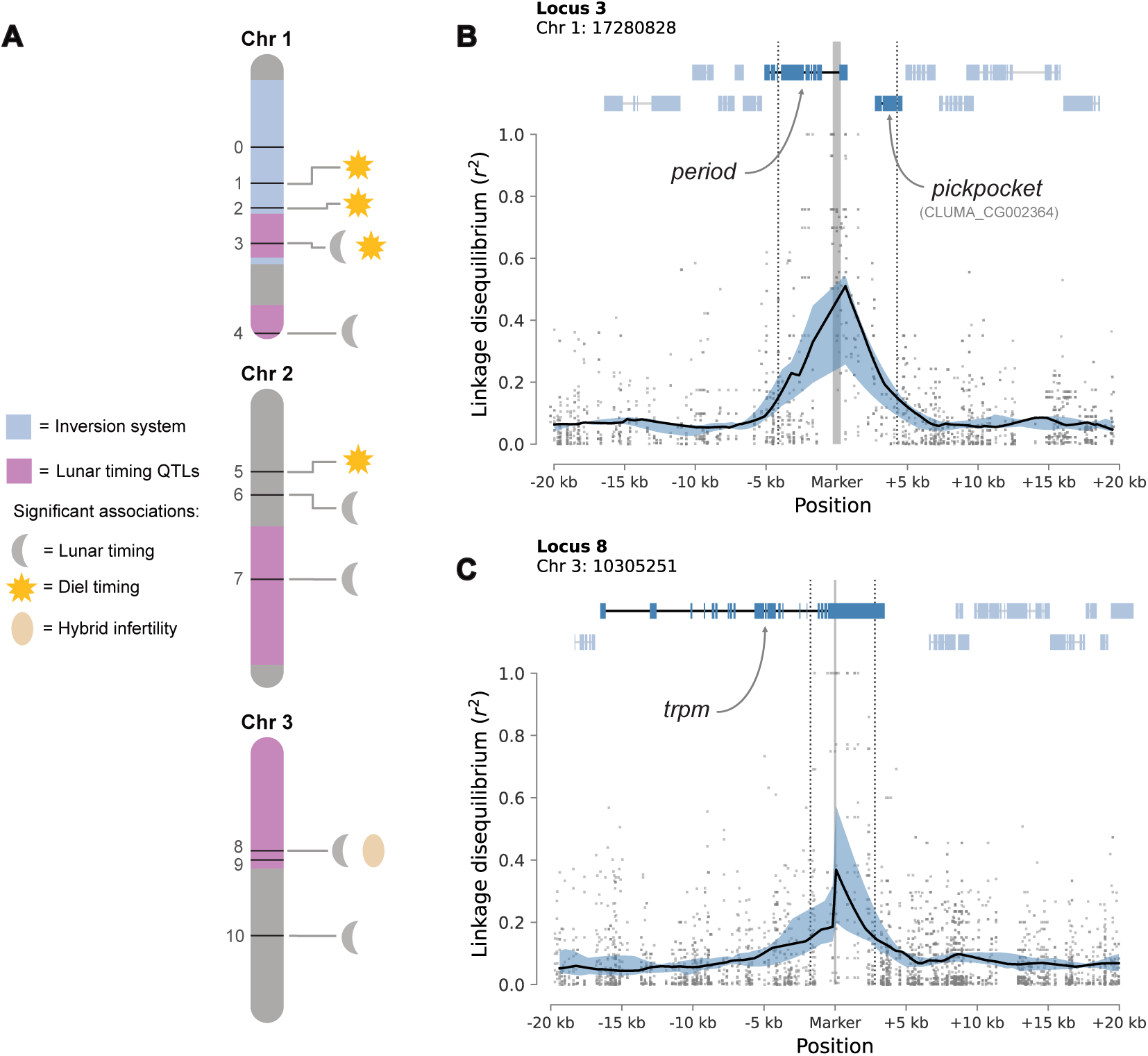
Association of genotypes with different isolation barriers. A) The high-F_ST_ loci (numbered 0-10) genotyped in the field couples for our study are associated with different barriers, giving an indication for the genomic basis for these barrier traits. Two of the loci (3 and 8) were associated with multiple barriers. The locations of the large inversion (1A) and lunar timing QTLs from (Briševac et al., 2023) are shaded in blue and pink, respectively. B) By visualizing the physical LD block around locus 3, we find that there are two genes at this locus – *period* and *pickpocket* – that are in tight linkage (dashed grey lines indicate a median LD > 0.2). C) The LD block surrounding locus 8 is entirely within the *trpm* gene. For B and C, LD was calculated for five proxy SNPs surrounding the marker SNP within each chronotype. A LOESS was fit to the LD values for each chronotype, and the median value of these LOESS’ were plotted as a solid black line. The upper and lower quartiles are represented by blue shading. Pairwise LD values between each proxy SNP and all other SNPs in the window are plotted as grey dots.

**Table 2.** Association testing between genotypes at the 11 high-F_st_ loci used in this study and the barriers to reproduction found to be acting in the field caught couples.

| **Locus** | **Lunar**<br>**timing** | **Diel timing** | **Hyrbid**<br>**infertility** |
| --- | --- | --- | --- |
| **0** | N.S. | N.S. | N.S. |
| **1** | N.S. | $\beta = -0.24$<br>$p = 2.10e-3$ | N.S. |
| **2** | N.S. | $\beta = 0.22$<br>$p = 3.43e-3$ | N.S. |
| **3** | $\beta = 0.24$<br>$p = 2.18e-3$ | $\beta = 0.73$<br>$p = 7.41e-25$ | N.S. |
| **4** | $\beta = -0.19$<br>$p = 2.15e-2$ | N.S. | N.S. |
| **5** | N.S. | $\beta = 0.19$<br>$p = 1.03e-2$ | N.S. |
| **6** | $\beta = -0.50$<br>$p = 1.59e-7$ | N.S. | N.S. |
| **7** | $\beta = -0.84$<br>$p = 1.74e-24$ | N.S. | N.S. |
| **8** | $\beta = -0.34$<br>$p = 3.82e-4$ | N.S. | $\beta = -1.00$<br>$p = 2.57e-2$ |
| **9** | N.S. | N.S. | N.S. |
| **10** | $\beta = -0.26$<br>$p = 4.86e-5$ | N.S. | N.S. |

Notably, two loci (3 and 8) were associated with multiple reproductive barrier traits (Table2). Locus 3, harboring the circadian clock gene *period* and a *pickpocket* gene (CLUMA_CG002364) encoding an ion channel, was associated with diel and lunar timing. Locus 8 overlaps with the *trpm* gene, which again encodes an ion channel, and is associated with lunar timing and hybrid infertility. Such multi-trait associations could arise either through pleiotropic effects of a single gene within a locus, or through tight physical linkage among multiple genes, each contributing to a different trait. To evaluate these alternatives, we examined patterns of physical linkage disequilibrium (LD) surrounding each locus (Figure 4B, C). At locus 3, the region of elevated LD surrounding the marker encompassed both genes, allowing for both linkage or pleiotropy. In contrast, at locus 8, elevated LD was confined entirely within the *trpm* gene, consistent with pleiotropic effects on both lunar timing and hybrid infertility.

## 5 Discussion

### 5.1 Lunar timing as a magic trait

Magic traits have long tantalized evolutionary biologists as they provide a simple mechanism through which speciation with gene flow is possible (Servedio et al., 2011). Of these traits, one that might be particularly important due to its commonality (Raible et al., 2017) and potential for driving speciation (Servedio et al., 2011) is divergent reproductive timing. Here, we investigated whether reproductive timing acted as a magic trait in the divergence of sympatric C. marinus chronotypes.

Adult C. marinus are restricted to mating at the waterline during low tides, and the height of low tide varies periodically over the lunar month. Therefore, the lunar day of emergence sets the depth of oviposition, and consequently where offspring live in the intertidal zone (Ekrem et al., 2025). This relationship implies that divergent selection on the depth at which larvae live also selects for divergence in lunar reproductive timing (Jacobsen et al., 2026). A major assumption of this hypothesis, however, is that lunar reproductive timing effectively isolates reproduction in the wild. Whole-genome sequencing of 24 individuals from each strain suggested there are genetically differentiated loci, though none of them was genetically fixed (Figure 1B; Brisevac et al 2023). Studies with larger numbers of individuals either defined the chronotypes solely based on phenotypes (Ekrem et al., 2025) or were based on laboratory strains (Kaiser et al., 2021). Here we genotyped 797 individuals caught in the field over the course of two lunar months. We found that individuals emerging in the full moon vs. the new moon peak indeed constitute largely distinct genetic groups, strongly supporting the hypothesis that lunar reproductive timing acted as a magic trait.

We quantified how effective lunar timing is at restricting gene flow and estimated that it has an RI value between 0.88 and 0.90, depending on the cross direction and dataset (field vs laboratory). This underlines that magic traits can play a pivotal role in speciation with gene flow, though it remains to be seen how commonly this is the case for other organisms. Despite their potential importance in driving speciation, estimates of RI for magic traits are still rare in the literature. RI due to assortative mating on wing color patterns in *Heliconius* species, a well-studied example of a magic trait (Kronforst et al., 2006; Smadja & Butlin, 2011), has been estimated to range from 0.07 to 0.98 depending on the species pair and crossing direction (Mérot et al., 2017).

Diet-based assortative mating in the cotton mouse *Peromyscus gossypinus* was estimated to be 0.48 (Delaney & Hoekstra, 2019), and size-based assortative mating in *Neodiprion* sawfly species was found to have an RI between 0.19 and 0.77 depending on the species pair (Glover et al., 2023). Although estimates of RI are sensitive to methodological and biological context, lunar timing in the Roscoff chronotypes nonetheless appears to fall toward the upper end of reported values for magic traits.

### 5.2 Strong RI despite minimal divergence

While lunar timing provided significant RI between chronotypes, it was not the only barrier to reproduction that we found. Diel timing, mate preference, and hybrid infertility all appear to be restricting gene flow. While these subsequent barriers only contribute marginally to the total RI as compared to lunar timing, their combined effect produces a total RI value between 0.93 and 0.94, again dependent on crossing direction and data set. This level of total RI is comparable with several examples of “good” species (e.g. 0.94 between *Argyroderma deleatii* ‘late flowering’ and *A. framesii* subsp. *framesii* (Boucher et al., 2024) or 0.895 for *Habenaria limprichtii* and *H. davidii* (H.-P. Zhang et al., 2022)), or taxonomically defined subspecies (e.g. 0.90-0.94 for *Heliconius melpomene aglaope* and *H. m. amaryllis* (Garzón-Orduña & Brower, 2018)).

Such strong RI between chronotypes was surprising as, unlike the examples above as well as many other species or ecotype pairs, there are no described morphological differences between chronotypes, and the only known ecological difference is the depth at which larvae live (Ekrem et al., 2025). Moreover, genetic divergence between chronotypes is low (F_ST_ = 0.012; Kaiser et al., 2021), with nearly all individuals sharing the same mitochondrial COI haplotype (Kaiser et al., 2021), consistent with a recent divergence that likely occurred within the last ∼10,000 years (Kaiser et al., 2010).

The rapid evolution of strong RI is not unprecedented; different *Clarkia* species were found to have evolved strong RI within 65,000 years (Briscoe Runquist et al., 2014) and freshwater/limnic species of sticklebacks also evolved strong RI within the last 13,000-15,000 years (Lackey & Boughman, 2017). However, the limited genetic, morphological, and potentially ecological differentiation accompanying strong RI is rare. One example are central American populations of *Sepsidae* dungflies (Puniamoorthy, 2014), which exhibit strong RI due to differences in courtship behavior but limited divergence in COI sequence and morphology. Another example are sympatric populations of *Nasonia vitripennis* which have evolved sexual and pre-mating barriers prior to ecological divergence (Malec et al., 2021). These examples, along with the Roscoff FM and NM chronotypes, potentially represent cases of the rapid evolution of strong RI in the face of gene flow, where fast-evolving mating traits drive speciation well ahead of neutral genetic, morphological, and perhaps ecological divergence. Further work characterizing the ecological niches of the sympatric *Clunio* chronotypes would help determine if this truly is the case.

### 5.3 Genomic mechanisms facilitating the buildup of RI

#### 5.3.1 Inversions

Several mechanisms are hypothesized to facilitate the rapid establishment of strong RI (Kulmuni et al., 2020). One that is relevant for the sympatric C. marinus chronotypes is chromosomal rearrangements. These rearrangements suppress recombination, allowing co-adaptive alleles at barrier loci to couple together (Berdan et al., 2023). A large inversion system on chromosome 1 was found to be differentiated between the Roscoff chronotypes, though there was no direct association between the inversions and lunar timing (Briševac, Peralta, et al., 2023). In our study, we found that this inversion system contained all but one of the diel timing associated loci we genotyped, as well as one locus associated with both diel and lunar timing (locus 3, containing the *period* gene). While this pattern of association may arise due to the correlation of genotypes (i.e. LD) within the inversion, we found that our genotypes exhibited no multicollinearity. Thus, the independent effects of these loci could be reliably estimated in our models, suggesting that the associations of several loci within the inversion with diel timing are not merely a statistical artifact. The inversion may play a larger role in diel differences between chronotypes than in lunar differences, potentially facilitating the coupling of these different diel timing loci, increasing the strength of the diel timing barrier, as well genetically coupling diel timing to lunar timing via locus 3.

#### 5.3.2 Polygenicity and physical linkage

Our results also point towards two other mechanisms relating to the genetic architecture of traits that might facilitate the rapid build-up of RI. The first is the increased chance of barrier gene co-localization for polygenic barrier traits. Since recombination opposes the coupling of barriers, and therefore the buildup of strong RI, genomic architectures which place barrier genes within close proximity (i.e. physical linkage) are more likely to result in speciation (Butlin & Smadja, 2018; Smadja & Butlin, 2011). If barrier traits are highly polygenic, like lunar timing may be for C. marinus (Fuhrmann et al., 2023), then there is an increased chance of co-localization of genes, meaning that it is more likely for two barrier genes to be in tight linkage (Servedio et al., 2011). This implies that polygenic traits under divergent selection are more likely to “catch” other barrier genes and couple with them than traits with more simple genomic architectures.

This might have been the case for locus 3 in our study, as two genes, *period* and *pickpocket*, were found to be within physical linkage of each other in a locus associated with both diel and lunar timing. *Period* is a core circadian clock gene across nearly all bilaterian clades (Stanton et al., 2022), and was previously found to be strongly differentiated between the two chronotypes and inside a quantitative trait locus (QTL) for lunar timing differences (Briševac, Peralta, et al., 2023). Given it’s well understood and conserved role in the circadian clock, it is likely that *period* could be involved the diel timing differences between chronotypes. As the circadian clock is known to play a crucial role in lunar time-keeping in C. marinus (D. Neumann, 1989; J. Neumann et al., 2024), it is also conceivable that mutations in period will also affect lunar timing via *modular pleiotropy* (i.e the involvement of the entire “circadian clock module” in lunar timing, see Bradshaw & Holzapfel, 2010). On the other hand, *pickpocket* is known from *Drosophila* to play a role in environmental sensing, and when knocked down, induces changes in the timing of behavioral transitions between the foraging and wandering larval stages (Ainsley et al., 2008). Lunar timing in C. marinus is known to involve sensing environmental cues for entrainment (i.e. synchronization) of the circa-lunar clock (Neumann 2014; Briševac, Prakash, et al., 2023), as well as a timed developmental arrest during the fourth larval instar synchronizes development and subsequent emergence (D. Neumann & Spindler, 1991). If *pickpocket* was to play a similar role in environmental sensing and developmental transitions in C. marinus as it does for *D. melanogaster*, then it may be responsible for the lunar timing association at locus 3. Functional validation of both *pickpocket* and *period* in C. marinus would help determine if this is the case.

#### 5.3.3 Pleiotropy and complex traits

The second mechanism suggested by our results is an increased chance of divergent selection acting on pleiotropic genes for complex barrier traits. The degree of pleiotropy associated with loci is known to vary, with mutations in genes or regulatory regions involved with many core biological processes likely affecting multiple traits (Mensch et al., 2008; J. Zhang, 2023). In general, it seems that the degree of pleiotropy for genes underlying complex traits is greater than that for Mendelian traits (Barbitoff et al., 2024), especially for genes involved in general biological functions (Watanabe et al., 2019). Therefore, divergent selection acting on traits that involve core biological processes, such as the timed developmental transition in C. marinus that gives rise to timed reproduction, would be more likely to recruit pleiotropic genes that could generate additional barriers, and thus stronger RI.

Locus 8 in our study might be an example of this, as it was associated with both lunar timing and hybrid infertility. The LD block surrounding the genotyped marker was entirely within the *trpm* gene, suggesting that this may be a case of pleiotropy. TRP channels are known to be polymodal environmental sensors, reacting to stimuli such as temperature and light (Diver et al., 2022; Zheng, 2013). Additionally, *trpm* has been found to be highly expressed in the gonads of planarian flatworms, where knockdowns indicated a role in spermatogenesis (Curry et al., 2025), as well as the eggs of the fall armyworm *Spodoptera frugiperda*, where it is suggested to play a role in Zn+ and Mg+ homeostasis (Su et al., 2025). Allelic variation in *trpm* for either FM or NM might therefore effect how larvae perceive lunar entrainment cues, while heterozygosity of these alleles might lead to the disruption of egg development, potentially through ionic disregulation. Again, direct functional validation of *trpm* in the FM and NM chronotypes would be required to determine if this is the case, and if the multiple effects of locus 8 are truly the product of pleiotropy.

### 5.4 Conclusions

Altogether, our results suggest that the rapid buildup of RI in the divergence of the FM and NM chronotypes involved several different mechanisms acting in synergy. First, divergent lunar reproductive timing–probably caused indirectly by selection on the depth of the larval habitat (Jacobsen et al., 2026)–acted as a magic trait and facilitated divergence in the face of gene flow. Secondly, the large inversion may have promoted the coupling of multiple barrier loci, especially those involved in diel timing. Finally, potentially due to the polygenic and complex nature of lunar timing, divergence at some loci was accompanied by the formation of additional barriers to gene flow via pleiotropy, and/or very tight linkage between barrier genes, resulting in the rapid evolution of strong RI.

While the hypotheses generated from our results are exciting, several limitations of our study warrant careful consideration. Specifically, our study focused only on genotypes from 12 high-F_ST_ loci, and therefore our association analysis, while powerful to detect associations at these 12 loci, did not consider the genomic background. Consequently, the locus–phenotype associations reported here should be interpreted as candidate relationships that motivate future functional and genomic validation, rather than as definitive causal links. At the same time, the exceptionally strong reproductive isolation between chronotypes, combined with their minimal morphological and genomic divergence, makes C. marinus a powerful natural system for dissecting the earliest stages of speciation.

