## Supplemental materials for "Lunar Reproductive Timing Acts as a Magic Trait and May Recruit Additional Isolating Barriers in Sympatric Marine Midge Populations"

Supplement

Figure S1

Lunar (A) and Diel (B) emergence timing of FM and NM chronotypes under laboratory conditions. The data is from (Kaiser et al., 2021)

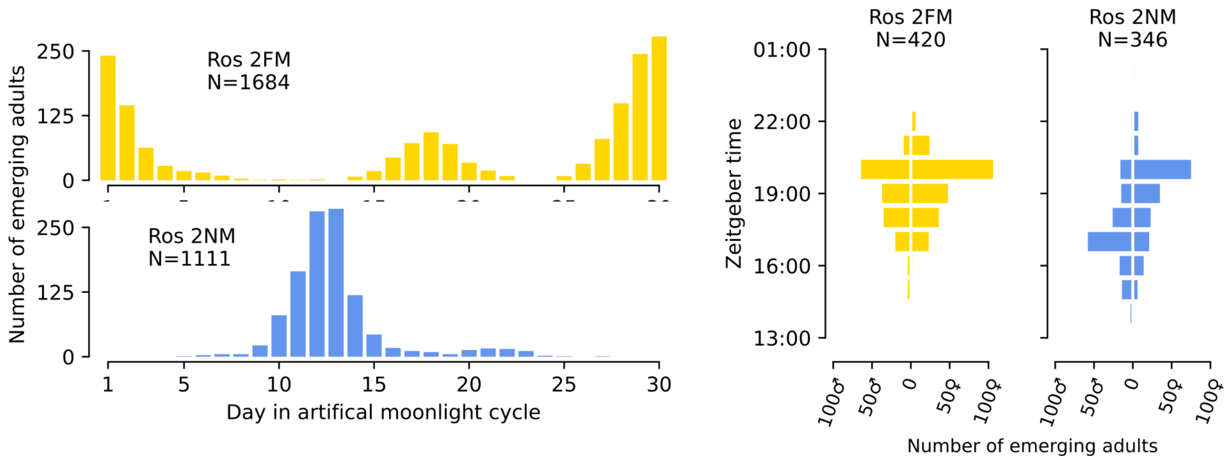

Figure S2

The fertilization rate (A) and hatching rate (B) for different crossing direction in the laboratory. Significant differences were tested using a Kruskal-Wallis test followed by a *post-hoc* Dunn test (p<0.05: *, p<0.001: **). A decrease in fertilization rate for conspecific crosses would indicate gametic incompatibility, while a decrease in hating rate would suggest hybrid depression

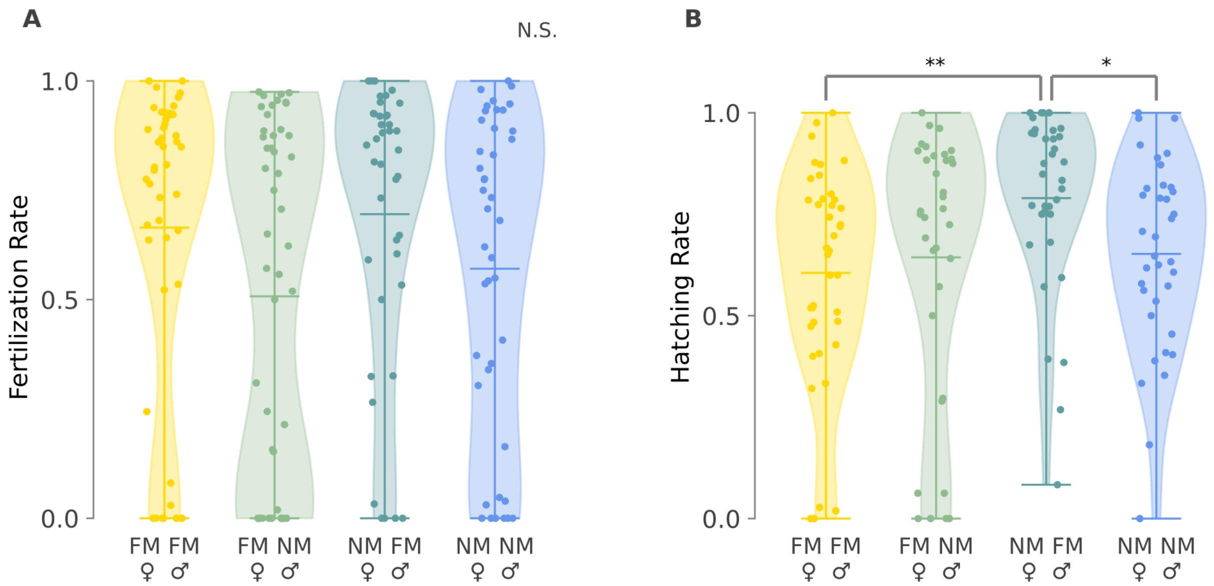

Figure S3

Fertilization rate (A) and offspring survival rate (B) for couples caught in the field. A negative effect of ancestry difference on fertilization rate would signify gametic incompatability, while a negative effect of expected heterozygosity in the offspring on their survival rate would indicate hybrid depression. Both were modeled via a Zero-altered negative binomial model and no significant effects were found.

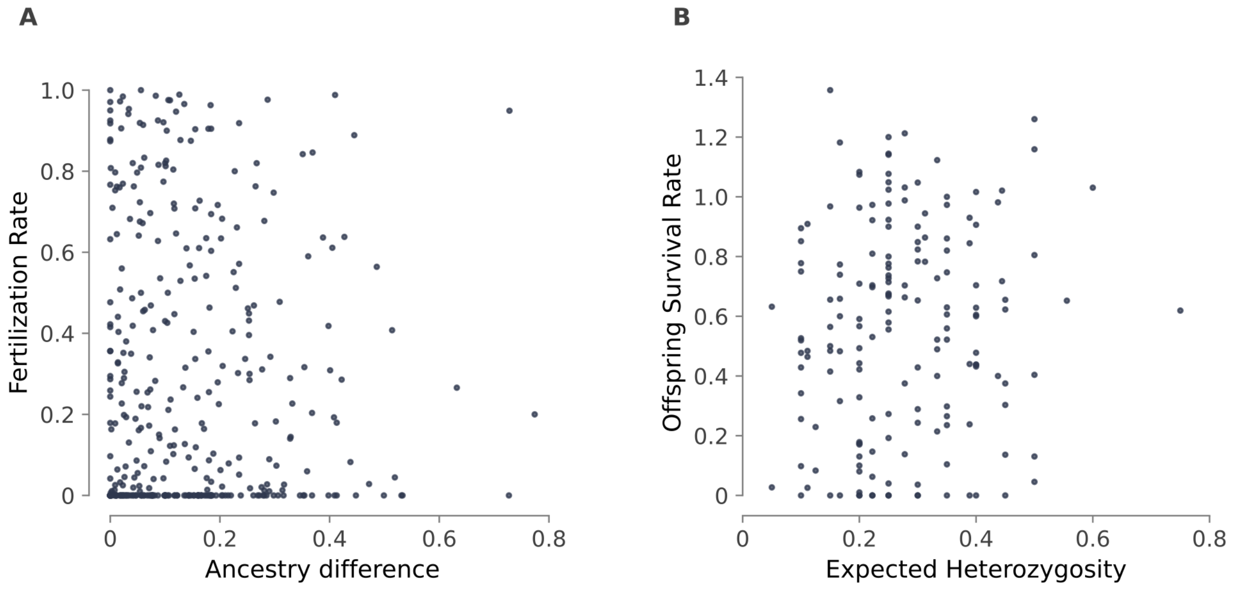

Figure S4

A) The strength of RI for mate choice depends on the age (minutes after emergence) of the female in a copula. We estimated the average age for females in each crossing direction by simulating couples based on the distribution of diel emergence timing for each chronotype in the lab. Since *C. marinus* individuals only live for a few hours, we censored any couples for which individuals were above a certain age threshold. As, the choice of this age threshold was somewhat arbitrary, we performed a sensitivity analysis to see how our choice for maximum age effects (B) the average female age for couples, (C) strength of RI for mate preference, and (D) the relative contribution of mate preference towards the total RI between chronotypes. All calculations were done for conspecific crosses, as well as the FM♀-NM♂ and NM♀-FM♂ directions.

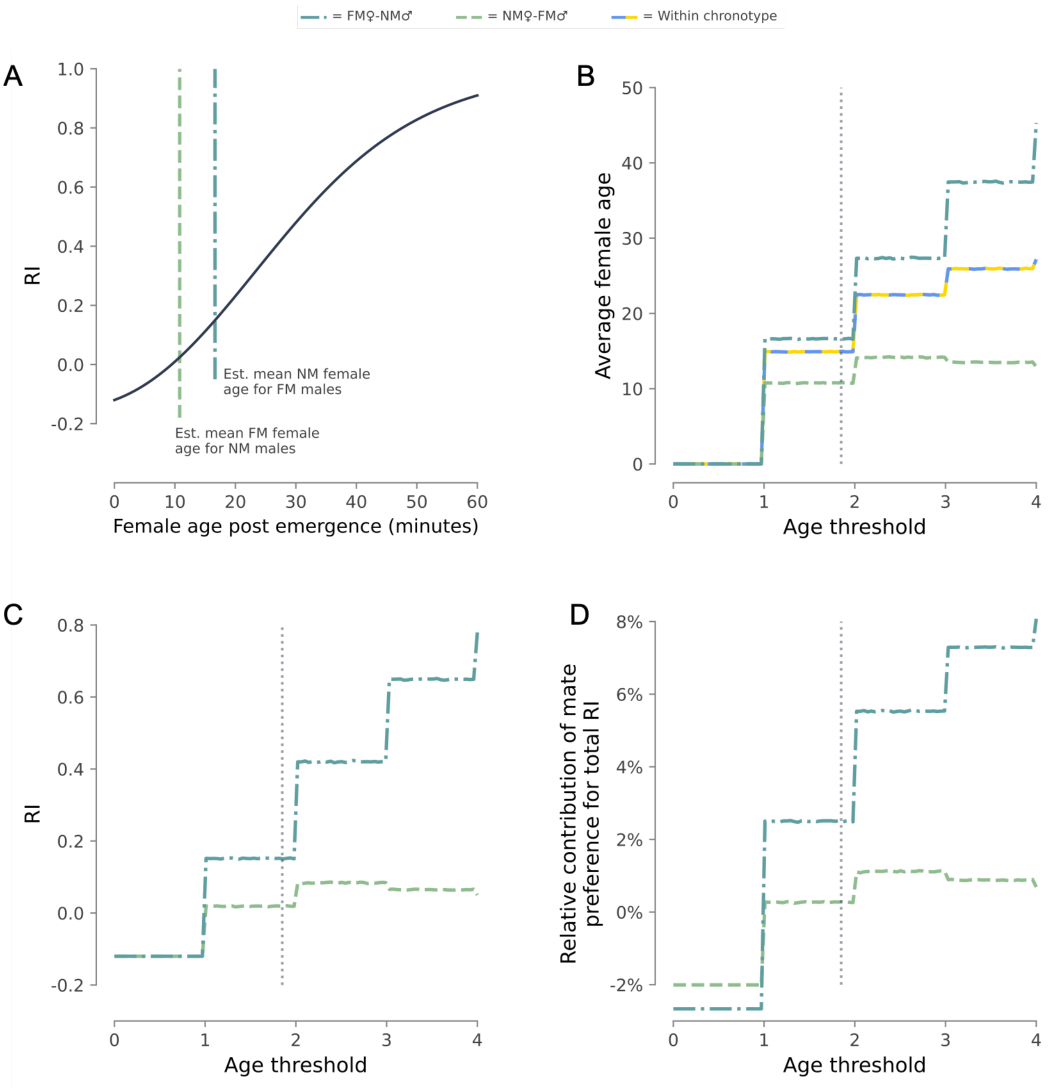

Table S1

Primers used for amplification of the genotyped loci.

| Locus | Chr | start | end | Forward Primer Sequence | Reverse Primer Sequence |
|---|---|---|---|---|---|
| Multiplex 1 | | | | | |
| 2 | 1 | 11755069 | 11755458 | TTAATAACACCTGCTTTGCG | AACGAGTTTAATTAGAAATTTAAAC |
| 3 | 1 | 14034017 | 14034145 | GTAGGTAGTTGAAGGTAAATTCTA | GAAATGCTCGACCACATCT |
| 4 | 1 | 17280400 | 17280648 | TGGAAAGGAAGCGGTAAAGA | AACAAACTCCCACGCAAAAA |
| 5 | 1 | 25531981 | 25532483 | TACTCATGAGGTGTTTCATTAAATTT | TGTGCATATATGTATGCTTTAACATC |
| 8 | 2 | 17316253 | 17316917 | TGTTGCAATTTGCTCACG | CGAGATTGCTGTAAGTTATTTTTA |
| 11 | 3 | 18085772 | 18086082 | AAAAAAAGAAGGTTCATATCTAAAT | CTGTAGGTTGCCTAGGAG |
| Multiplex 2 | | | | | |
| 1 | 1 | 8454348 | 8454703 | AACAGGTTTTTGTTCGGTGA | CCAACATCTGAGCCAGCTCTC |
| 6 | 2 | 7465168 | 7465728 | GGAAGCCAATCAGTGGAAACC | CCACATTTTCCCATCAATTCG |
| 7 | 2 | 9578462 | 9578670 | GTGGTCGTCCGCCTTTAGAT | TACTTGATGACGCGGTGGTT |
| 9 | 3 | 10304898 | 10305352 | AGGCTCGTCCGTGACGTATT | CCTTCGACACATTCCTCTTCG |
| 10 | 3 | 11136148 | 11136435 | TTTTTGGTTGGGGAGCTTCT | CTGATTCTGATGCACAATTGGA |
| Insertion-deletion (indel) in the *period* locus | | | | | |
| *per* | 1 | 17280546 | 17281110 | ACAACGTGACCTGTGACAAT | GAATACTGAGTGTAAGACTTGGC |

Table S2

The absolute values of RI for different reproductive barriers, calculated from both laboratory and field data and for both crossing directions.

| Barrier | Measurement | Direction | Absolute RI | 95% CI |
|---|---|---|---|---|
| Circalunar timing | Field | FM♀-NM♂ | 0.88 | 0.83-0.93 |
| | | NM♀-FM♂ | 0.90 | 0.86-0.95 |
| | Laboratory | Either | 0.90 | 0.89-0.92 |
| Circadian timing | Field | FM♀-NM♂ | 0.78 | 0.73-0.86 |
| | | NM♀-FM♂ | 0.80 | 0.75-0.86 |
| | Laboratory | FM♀-NM♂ | 0.52 | 0.45-0.61 |
| | | NM♀-FM♂ | 0.13 | 0.10-0.25 |
| Mate preference | Laboratory | FM♀-NM♂ | 0.02 | -0.11-0.16 |
| | | NM♀-FM♂ | 0.15 | -0.01-0.38 |
| Hybrid infertility | Field | Either | 0.25 | 0.12-0.39 |

Table S3

The values of total RI between chronotypes along the sequence of reproductive barriers. Again, calculations were performed independently for data measured from the field or laboratory couples, as well as for both crossing directions.

| Barriers | Direction | Value | 95% CI |
|---|---|---|---|
| Field measurements | | | |
| Lunar timing | FM♀-NM♂ | 0.88 | 0.83-0.93 |
| | NM♀-FM♂ | 0.90 | 0.86-0.95 |
| Lunar timing + Diel timing | FM♀-NM♂ | 0.89 | 0.86-0.95 |
| | NM♀-FM♂ | 0.92 | 0.89-0.97 |
| Lunar timing + Diel timing + Hybrid infertility | FM♀-NM♂ | 0.94 | 0.91-0.97 |
| | NM♀-FM♂ | 0.95 | 0.94-0.98 |
| Laboratory measurements | | | |
| Lunar timing | FM♀-NM♂ | 0.90 | 0.89-0.92 |
| | NM♀-FM♂ | 0.90 | 0.89-0.92 |
| Lunar timing + Diel timing | FM♀-NM♂ | 0.93 | 0.92-0.94 |
| | NM♀-FM♂ | 0.91 | 0.90-0.93 |
| Lunar timing + Diel timing + Mate preference | FM♀-NM♂ | 0.93 | 0.92-0.95 |
| | NM♀-FM♂ | 0.93 | 0.91-0.96 |
